# Cortical changes in motor preparation facilitate changes in motor execution following an intervention in severe chronic hemiparetic stroke

**DOI:** 10.1101/547083

**Authors:** Kevin B. Wilkins, Julius P.A. Dewald, Jun Yao

## Abstract

Effective interventions have demonstrated the ability to improve motor function by reengaging ipsilesional resources, which has been shown to be critical and feasible for hand function recovery even in individuals with severe chronic stroke. However, previous studies focus on changes in brain activity related to motor execution. How changes in motor preparation may facilitate these changes at motor execution is still unclear. In order to address this question, we had 8 individuals with severe chronic hemiparetic stroke participate in a device-assisted intervention. We then quantified changes in both coupling between regions during motor preparation and changes in topographical cortical activity at motor execution for both the hand in isolation and together with the shoulder using high density EEG. We hypothesized that intervention-induced changes in cortico-cortico interactions during motor preparation would lead to changes in activity at motor execution specifically towards an increased reliance on the ipsilesional hemisphere. We found that, following the intervention, individuals displayed a reduction in coupling from ipsilesional M1 to contralesional M1 within gamma frequencies during motor preparation for hand opening. This was followed by a reduction in activity in the contralesional primary sensorimotor cortex during motor execution. Meanwhile, during lifting and opening, a shift to inhibitory coupling within ipsilesional M1 from gamma to beta frequencies was accompanied by an increase in ipsilesional primary sensorimotor cortex activity following the intervention. Together, these results show that intervention-induced changes in coupling within or between motor regions during motor preparation lead to corresponding changes in focal cortical activity at execution.

## 1. Introduction

In individuals with stroke, improvements following an intervention are at least partially mediated by neural changes. One of the more common findings for effective interventions is a return to “normal” cortical activity patterns characterized by an increased reliance on the ipsilesional hemisphere that resembles patterns observed in healthy controls^1,2^. To this point, most investigations of intervention-induced neural changes have focused on cortical activity related to motor execution ^3–5^. However, the proper motor command during execution requires appropriate motor preparation. How intervention-induced changes in motor preparation may facilitate cortical changes related to execution is still unknown.

In healthy controls, the temporal-spatial feature of motor preparation related to non-visually guided hand/finger movements is characterized by a flow of information from secondary motor areas to contralateral primary motor cortex^6–8^. In addition, previous results reported decreased power (i.e., desynchronization) in beta (13-30 Hz) and mu (8-12 Hz) during motor preparation, and an association between such desynchronization and the release of the motor command^12,13^. Meanwhile, power in higher gamma frequencies (30-80 Hz) increases (i.e., synchronization) during movement preparation. Unlike lower frequencies, this increase in gamma reflects local intracortical processing rather than descending motor commands^14,15^.

Following a stroke, spatial and frequency features during motor preparation have been shown to be altered. For example, previous studies have reported overactivation of the supplementary motor area (SMA)^9^ as well as increased overlap of limb representations on the cortex^10^, which is especially pronounced in individuals with severe impairments. Post-stroke individuals also display stroke-induced changes in these neuronal oscillations during movement such as reduced beta desynchronization and changes in gamma-beta coupling between sensorimotor regions^16,17^.

Following interventions, changes occur not only at motor performance level but also in motor related brain activity. The majority of intervention studies to date have focused on cortical-changes related to motor execution. We argue that understanding how changes in motor preparation may facilitate changes in motor execution is also critical. One piece of the evidence in favor of this is from Norman and colleagues, who showed that specifically training preparation-related cortical oscillations led to subsequent improvements in motor function, further cementing the crucial role for motor preparation in proper movement^11^. Given the critical role of motor preparation in performance, investigating how these frequency characteristics during motor preparation change following an intervention would provide additional details into the nature of any observed cortical reorganization.

The goal of this study was to examine how cortical changes in motor preparation may facilitate changes in motor execution following an intervention that targeted hand/arm function recovery in individuals with severe chronic hemiparetic stroke. We hypothesized that intervention-induced changes in cortico-cortico interactions during motor preparation would lead to changes in activity at motor execution specifically towards an increased reliance on the ipsilesional hemisphere. To test this hypothesis, we examined dynamic cortical coupling between motor regions during motor preparation and cortical activity within motor regions at motor execution following an effective device-assisted intervention. Through this combined approach, we explored how the interactions between regions during motor preparation gave rise to any intervention-induced activity changes. Specifically, we investigated regions that synchronize power at the same frequency (i.e., linear coupling) or across different frequencies (i.e., nonlinear coupling) because the well-established physiological underpinnings of specific frequency bands within the motor system would provide insight into the underlying neural mechanisms that may be shaping cortical activity at motor execution^20–23^.

## 2. Methods

The EEG-data during the motor execution phase related to pure hand opening have been reported before^5^. Here, we analyzed these data, however, during motor preparation phase, to investigate the cortico-cortico interactions during motor preparation. Other data related to hand opening while lifting the arm are original.

### 2.1 Stroke participants

Eight individuals with chronic hemiparetic stroke (mean age: 63.5±4) with subcortical lesions and severe impairment (UEFMA: 11-24) participated in this study. Participant information and the methods of the intervention have been reported before^5,24^. We re-state this information here to make this paper self-contained. All individuals were screened for inclusion by a licensed physical therapist. Inclusion criteria included a UEFMA between 10 and 30 out of 66, no botulinum toxin within the last 6 months, MRI compatibility, the ability to elicit enough EMG activity at wrist/finger extensors, the ability for the FES to generate a hand opening of at least 4 cm between thumb and index finger, the ability to lift the paretic arm, and the ability to give informed consent. See Table 1 for full demographics. This study was approved by the Northwestern University institutional review board. All procedures performed in studies involving human participants were in accordance with the ethical standards of the institutional and/or national research committee with the 1964 Helsinki declaration and its later amendments or comparable ethical standards. All participants gave informed written consent.

**Table 1.** Participant demographics and clinical characteristics

| Participant | Age | Time Since<br>Stroke (yrs) | Affected<br>Hand | UE FMA | CMSA |
| --- | --- | --- | --- | --- | --- |
| P01 | 64 | 9 | R | 23 | 3 |
| P02 | 62 | 8 | L | 12 | 3 |
| P03 | 68 | 3 | L | 17 | 3 |
| P04 | 61 | 22 | L | 11 | 3 |
| P05 | 61 | 13 | L | 24 | 3 |
| P06 | 70 | 20 | R | 13 | 3 |
| P07 | 59 | 6 | R | 24 | 3 |
| P08 | 63 | 9 | R | 22 | 4 |

Note: Upper Extremity Fugl Meyer Assessment (UE FMA); Chedoke McMaster Stroke Assessment (CMSA)

### 2.2 Intervention Design

Individuals took part in a 7-week intervention with 3 sessions per week lasting approximately 2 hours. During each session, individuals completed 20-30 trials of a reach-grasp-retrieve-release task with a jar while seated at a table. A novel EMG-FES device, called ReIn-Hand^25^, was used to assist paretic hand opening since these individuals lacked sufficient finger extension ability to complete the task without the stimulation. The novelty of this device is that it can detect hand opening intent and drive the paretic hand open even during lifting and reaching movements that typically increase expression of the flexion synergy at the elbow^26^, wrist, and fingers^27,28^. Briefly, this device recorded EMG activity from eight muscles (deltoid, biceps brachii, triceps, extensor communis digitorum, extensor carpi radialis (ECR), flexor digitorum profundus, flexor carpi radialis (FCR), and abductor pollicis), and used EMG features to detect hand opening intent in real-time to trigger an Empi transcutaneous electrical neuro-stimulation device (Vista, CA, USA). The stimulation electrodes were applied to the wrist/finger extensors with the following settings: biphasic symmetrical waveform, frequency = 50 Hz ± 20%, pulse width = 300 μs, amplitude = sufficient for maximum hand opening without discomfort, and duration = 3 s. The ReIn-Hand device allows for intuitive control since the same muscles the participant is attempting to drive (i.e. finger/wrist extensors) are being used to control the stimulation. All participants could successfully use the device to complete the described task.

### 2.3 EEG Experiment and Behavioral Task

Participants took part in an EEG experiment within one week prior to and following the intervention. Participants sat in a Biodex chair (Biodex Medical Systems, Shirley, NY) with straps crossing the chest and abdomen to limit potential trunk movements. The participant’s paretic arm was placed in a forearm-hand orthosis attached to the end effector of an admittance controlled robotic device (ACT^3D^) instrumented with a six degree of freedom (DOF) load cell (JR^3^ Inc., Woodland, CA). At the beginning of each trial, participants moved their hand to a home position, with the shoulder at 85° abduction, 40° flexion, and the elbow at 90° flexion angle. The participant received an auditory cue once they reached the home position. Following the cue, the participant relaxed at the home position for 5-7 s and then self-initiated one of 2 movements: 1) maximum paretic hand opening with the arm resting on a haptic table, or 2) maximum paretic hand opening while lifting against 50% of their maximum shoulder abduction (SABD) force. This shoulder abduction level was used since it is roughly equivalent to the weight of the limb in moderate to severe chronic stroke participants, thus making it a functionally relevant shoulder abduction level for translation to many activities of daily living that require simultaneously using the hand while lifting the arm. Participants were instructed to avoid eye movements by focusing on a point and movements of other body parts during the performance of each trial, which was confirmed by electrooculogram (EOG) traces and visual inspection by the experimenter, respectively. Participants performed 60-70 trials of each condition, which were separated into blocks (one block consisted of 20-30 trials of a particular condition). Blocks were randomized to minimize any order effects. Rest periods varied between 15 to 60 seconds between trials and 10 minutes between blocks.

Scalp recordings were made with a 160-channel High Density EEG system using active electrodes (Biosemi, Inc., Active II, Amsterdam, The Netherlands) mounted on a stretchable fabric cap based on a 10/20 system. The impedance was kept below 50 kΩ for the duration of the experiment. Simultaneously, EMGs were recorded from the extensor carpi radialis, flexor carpi radialis, and deltoid of the paretic arm, which were used to detect movement onset for post-processing purposes. All data were sampled at 2048 Hz. Additionally, the positions of EEG electrodes on the participant’s scalp were recorded with respect to a coordinate system defined by the nasion and pre-auricular notches using a Polaris Krios handheld scanner and reflective markers (NDI, Ontario, Canada). This allowed for coregistration of EEG electrodes with each participant’s anatomical MRI data.

### 2.4 Structural Imaging of the Brain

Participants participated in MRI scans at Northwestern University’s Center for Translation Imaging on a 3 Tesla Siemens Prisma scanner with a 64-channel head coil. Structural T1-weighted scans were acquired using an MP-RAGE sequence (TR=2.3s, TE=2.94ms, FOV=256×256mm^2^) producing an isotropic voxel resolution of 1×1×1 mm^3^. Visual inspection of acquired images was performed immediately following the data acquisition to guarantee no artifacts and stable head position.

### 2.5 Data Analysis for Cortico-cortical Connectivity During Motor Preparation

To investigate the cortico-cortico connectivity, we used dynamic causal modeling for induced responses (DCM-IR)^29^ to model the task-related time-varying changes in power both within and across a range of frequencies by estimating the coupling parameters within and between sources in a network. This approach has been used in previous hand movement tasks to elucidate the dynamic interactions within a motor network^6–30,31^.

#### 2.5.1 Definition of model space

Our motor network model consisted of 5 ROIs, including bilateral primary motor cortex (M1), bilateral premotor cortex (PM), and supplementary motor area (SMA). Locations of each of these regions were adapted from the Human Motor Area Template (HMAT)^32^ and are shown in Table 2. Bilateral primary sensory cortices were not included to reduce the computational demand and complexity of the model. Bilateral SMAs were treated as a single source due to their mesial position on the cortices. SMA also served as the input to the modelled network. It was chosen due to its critical role in motor preparation during self-initiated motor tasks^33,34^, and has previously been demonstrated to be an appropriate input for self-initiated motor tasks using DCM-IR^6–30,31^.

**Table 2.**
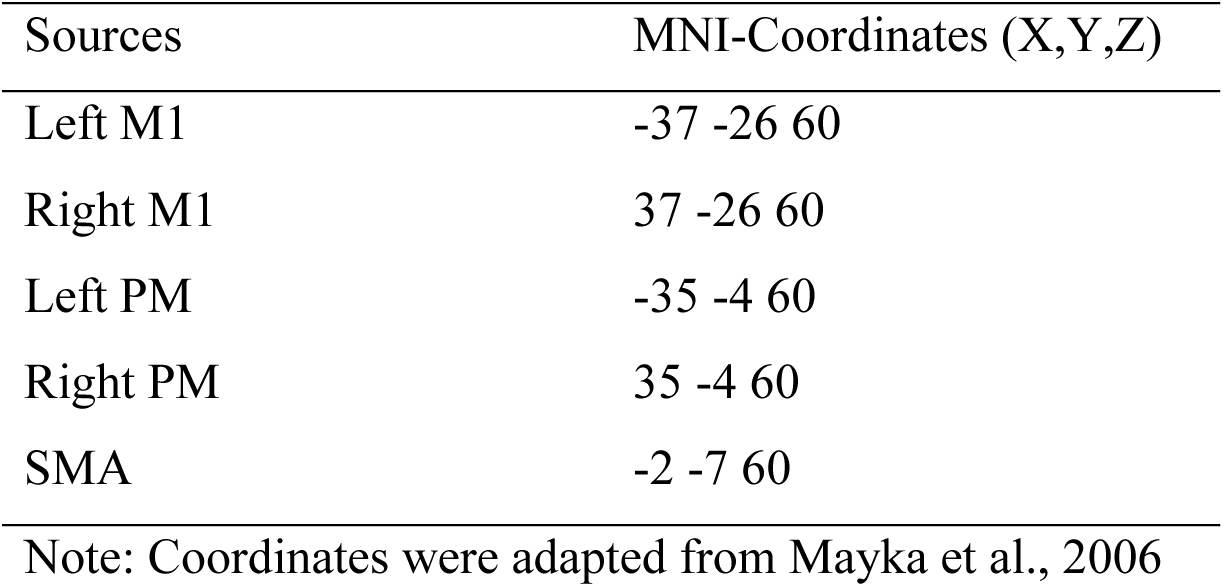
Coordinates of motor network

Different within- and cross-frequency connections between these 5 sources were used to create 12 models as shown in Supplementary Figure 1 which have successfully been used before in a similar motor task in healthy controls^30^. These 12 models were separated into 2 groups. Group 1 (models 1 to 6) allowed nonlinear and linear extrinsic (between region), but only linear intrinsic (within region) connections. Group 2 (models 7 to 12) allowed both nonlinear and linear connections for both extrinsic and intrinsic connections. Within each group, the 6 models consisted of 1 fully connected model, and the other 5 models missing 1 or 2 connections that were from one premotor area (PM) to either the other PM or to M1. Using this model, we tested the importance of nonlinear frequency interactions within regions as well as the importance of various connections to premotor regions.

#### 2.5.2 DCM Preprocessing

EEG data were preprocessed using SPM12 (SPM12, Wellcome Trust Centre for Neuroimaging, www.fil.ion.ucl.ac.uk/spm/). EEG was bandpass filtered between 1 and 50 Hz, segmented into trials (−2200 to +500 ms with 0 ms indicating EMG onset), and baseline-corrected. Trials exhibiting artifacts were removed. Artifact free trials were projected from channel space to the sources using the generalized inverse of the lead-field matrix using an equivalent current dipole for our chosen sources (see below)^29^. The spectrogram from 4 to 48 Hz at each source was computed using a Morlet wavelet transform (wavelet number: 7). This range includes theta (4-7 Hz), alpha (8-12 Hz), beta (13-30 Hz), and gamma (31-48 Hz) frequencies. The spectrogram was then averaged over all trials, cropped between −1000 to 0 ms, and then baseline-corrected by subtracting the frequency-specific power of the first 1/6 samples of the time window (−1000 to −833 ms). The input from SMA to the whole network was modelled as a gamma function with a peak at 400 ms prior to EMG onset with a dispersion of 400 ms. These values were chosen in order to capture the peak of the bereitschaftspotential during a self-initiated movement^35^. The model simulation was restricted to the time leading up to EMG onset (−1000 to 0 ms) to capture purely the motor preparation and command, rather than any potential sensory feedback related to the task. The dimensionality of the spectrogram was then reduced to four modes using singular value decomposition (SVD). The four modes preserved > 96% of the data variance on average. This dimensionality reduction both reduced the computational demand of the model inversion and denoised the data.

#### 2.5.3 Calculation of coupling parameters

For each of the models shown in Supplementary Figure 1, the dynamics of the spectrogram were evaluated using the following equation for each model described above:

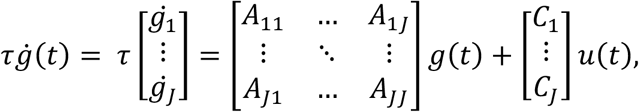

where vectors *g* and 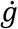 represent the instantaneous power and its first derivative at each of the modes (results of SVD) for each of the sources in the motor network. The A matrix contains the coupling parameters within and across different modes between any 2 regions within the *J*=5 regions, and the C matrix contains the weights of the extrinsic input 𝑢 from SMA. Each of the elements in the coupling A matrix refers to the influence of power at a specific frequency in one motor region on the power at another frequency in another region. Positive (i.e. excitatory) or negative (i.e. inhibitory) coupling suggests changes in power in the first frequency and region lead to the same directional or opposite change, respectively, in power in the second frequency and region. τ is a scaling factor and *t* represents time. Using the above equation and the output of the SVD, the DCM-IR method optimizes the A and C matrices to best describe the spectrogram of the measured data. The quality of a model and the estimated A and C matrices was quantified by the accounted variance from the predicted spectrogram.

#### 2.5.4 Bayesian model selection

We performed Bayesian model selection (BMS) with random effects^36^ on the 12 models described above for both the hand opening and the simultaneous lifting and opening conditions using the data from all of the participants to assess which model best explained the observed data. BMS with random effects was chosen since it is better equipped to handle potential heterogeneity associated with the study of a diseased population such as stroke^36^, and it contains a complexity term that penalizes a model based on the number of parameters it uses. The winning model, which was then used for further analysis of intervention-induced changes, was chosen based on the highest posterior exceedance probability (i.e. the probability that a given model is more likely than any of the other models tested).

#### 2.5.5 Inference on coupling parameters

Predicted spectra and A matrices from the four modes were projected back to frequency domain allowing for characterization of the coupling parameters as a function of frequency for the winning model. The coupling matrices for each intrinsic and extrinsic connections in the winning model for each participant were further smoothed with a Gaussian kernel (full-width half-maximum of 8 Hz) for each condition. These matrices include the frequency-to-frequency (both within- and cross-frequency) coupling values for each connection.

### 2.6 Data Analysis for Cortical Topography During Motor Execution

EEG data were aligned to the earliest EMG onset of the 3 muscles and segmented from −2200 to +200 ms (with EMG onset at 0 ms) using Brain Vision Analyzer 2 software (Brain Products, Gilching, Germany) and baseline-corrected (from −2180 to −2050 ms). Data were then visually inspected for the presence of artifacts. Trials exhibiting artifacts (e.g., eye blinks) were eliminated from further analysis. The remaining EEG trials were low pass filtered at 70 Hz and ensemble-averaged. The averaged EEG signals were down-sampled to 256 Hz and imported into CURRY 6 (Compumedics Neuroscan Ltd., El Paso, TX). The cortical current density strength (μA/mm^2^) from 150 ms to 100 ms prior to EMG onset was computed using the standardized low-resolution electromagnetic brain tomography (sLORETA) method (Lp = 1) based on a subject-specific boundary element method model with the regulation parameter automatically adjusted to achieve more than 99% variance accounted for. Possible sources were located on a cortical layer with 3 mm distance between each node. Although the inverse calculation was performed over the whole cortex, only the activity in the specific regions of interest (ROIs) was further analyzed. These ROIs included bilateral primary sensorimotor cortices (primary motor cortex (M1) + primary sensory cortex (S1)) and secondary motor cortices (supplementary motor area (SMA) + premotor area (PM)).

We quantified a cortical activation ratio 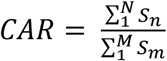 for each of the 4 ROIs, where s_n_ represents the current density strength of the n^th^ node, and n and m are the indices of nodes in a specific ROI and the whole sensorimotor cortices, respectively. The cortical activity ratio reflects the relative strength from one ROI as normalized by the total combined strength of the 4 ROIs.

### 2.7 Statistical Analysis for Cortical Activity

Statistics were performed using SPM and SPSS (IBM, V24). For the coupling parameters from the DCM analysis, T-statistics were used to calculate statistical parametric maps separately for each connection and condition pre/post intervention. Significance for intervention-induced changes in specific coupling parameters was set at *p* < 0.05 with family wise error (FWE) correction. For the cortical activity ratio during motor execution, a 2 (time) x 2 (task) x 4 (region) repeated measures ANOVA was performed after checking the data did not violate Mauchly’s sphericity test. We performed post-hoc paired t-tests when a main ANOVA effect or interaction was found. Significance was set a *p* < .05. Individual data are depicted for significant findings.

## 3. Results

ReIn-Hand assisted reaching-grasping training improved the score of Box and Block test significantly, as well as scores in other clinical assessments as reported previously^5,24^. The results included here will focus exclusively on the observed cortical neural changes pre-versus post-intervention.

### 3.1 Bayesian model selection and model fit

In order to determine any intervention-induced changes in connectivity, we first had to evaluate which DCM model tested best explained the observed data. BMS with random effects clearly preferred model 12 for each condition (see Supplementary Table 1), which had full connections between the 5 motor regions of interest and allowed both within- and cross-frequency interactions for intrinsic and extrinsic connections. Exceedance probabilities were .973 for the opening condition and .975 for the simultaneous lifting and opening condition. Supplementary Figure 2 depicts both the observed and predicted spectrograms in each of the 5 motor regions using the winning model for one participant during the hand opening condition. Comparison of these two spectrograms shows the overall similarity between the observed data (i.e., power changes over time) and the model-predicted data. Overall, this model explained ~85% of the original spectral variance for each condition, indicating that it was suitable for evaluating any intervention-induced changes.

### 3.2 Intervention-induced changes in coupling parameters

Once we determined the model that best explained the observed data, we examined whether any intervention-induced changes in connectivity for any of the region-region connections during motor preparation occurred. We found that the intervention induced two significant changes in the coupling parameters, one for each motor task. After the intervention, participants demonstrated significantly less excitatory coupling from ipsilesional M1 to contralesional M1 in gamma frequencies (47 Hz → 36 Hz) during hand opening (see Figure 1A). When looking at the individual coupling values for this particular regional coupling, we found that prior to the intervention, 6 out of 8 participants demonstrated positive coupling values, indicating that increases in gamma in ipsilesional M1 led to increases in gamma in contralesional M1 (see Figure 1B). However, following the intervention, 5 out of 8 participants showed zero or negative coupling, indicating that increases in gamma in ipsilesional M1 no longer led to increases in gamma in contralesional M1 (see Figure 1B).

**Figure 1.**
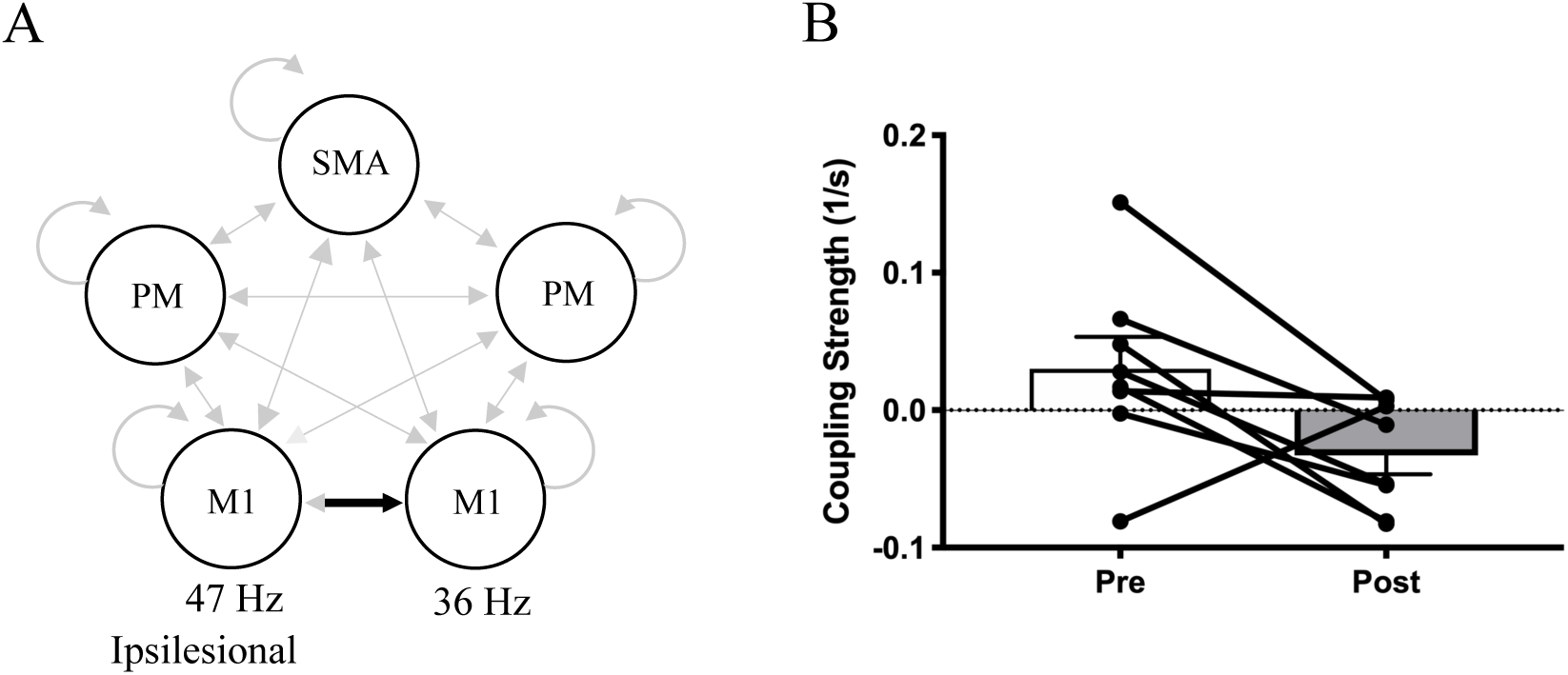
Reduced interhemispheric coupling following the intervention for hand opening on the table during motor preparation. (A) Schematic of significant decreases in coupling (black) from ipsilesional M1 (47 Hz) to contralesional M1 (36 Hz) within the motor network (light gray) following the intervention. None of the other region interactions (light gray arrows) showed significant changes pre to post intervention. (B) Average coupling strength data with individual data overlaid pre/post intervention for the connection depicted in A. 6 out of 8 subjects demonstrate a reduction in coupling strength for this M1-M1 connection within gamma frequencies. Positive values indicate excitatory coupling, while negative values indicate inhibitory coupling. Error bars represent SEM.

For the task of simultaneous lifting and opening, the intervention induced significantly more negative or inhibitory coupling within ipsilesional M1 from gamma to beta (44 Hz → 25 Hz) (see Figure 2A). When looking at the individual coupling values for this particular regional coupling, we found that prior to the intervention, 6 out of 8 participants showed positive coupling values, indicating that increases in gamma power within ipsilesional M1 led to subsequent increases in beta power also within ipsilesional M1 (see Figure 2B). However, following the intervention, 7 out of 8 participants showed negative or inhibitory coupling, indicating that increases in gamma power within ipsilesional M1 led to subsequent decreases in beta power within ipsilesional M1 (see Figure 2B).

**Figure 2.**
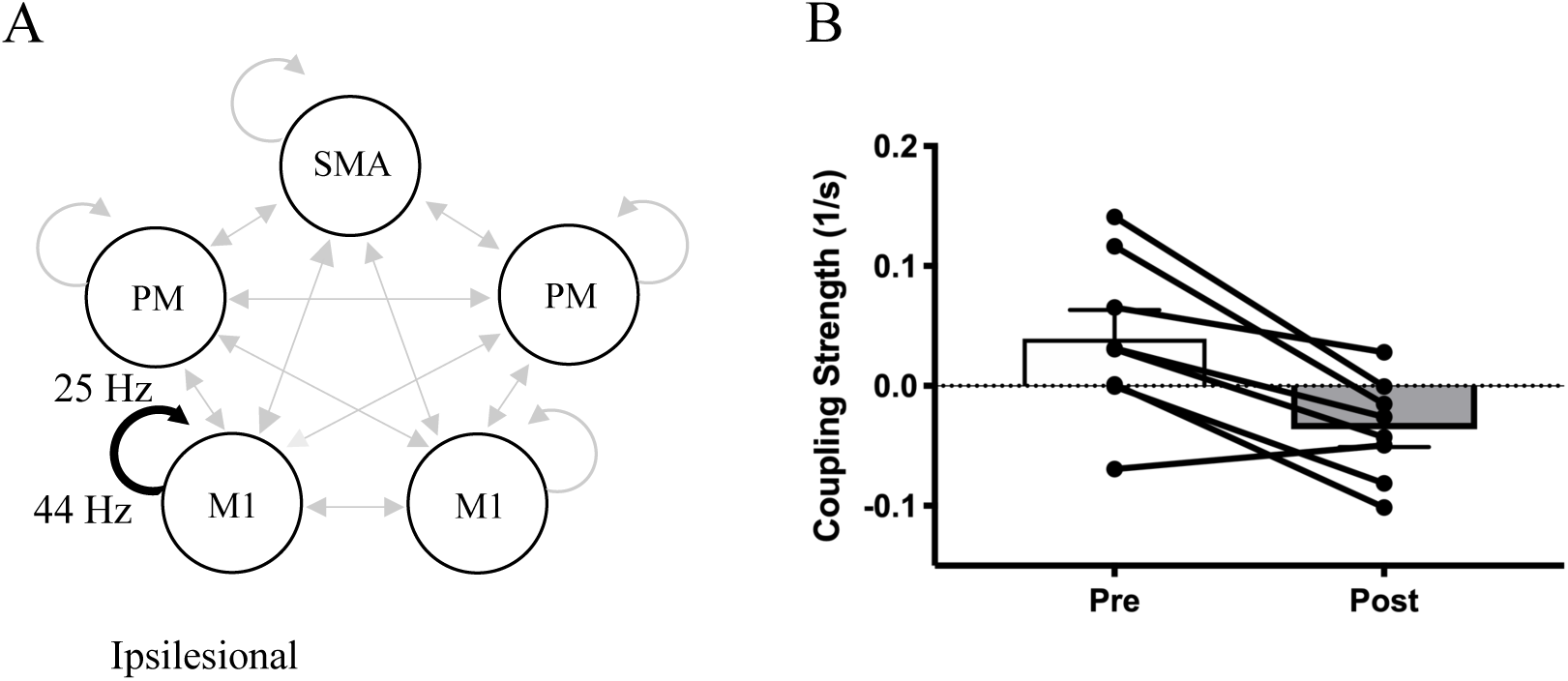
Altered intrinsic coupling within ipsilesional motor cortex following the intervention for hand opening while lifting during motor preparation. (A) Schematic of significant decreases in intrinsic coupling (black) in ipsilesional M1 (44 Hz to 25 Hz) within the motor network (light gray) following the intervention. None of the other region interactions (light gray arrows) showed significant changes pre to post intervention. (B) Average coupling strength data with individual data overlaid pre/post intervention for the connection depicted in A. 7 out of 8 subjects demonstrate negative coupling within ipsilesional M1 from gamma to beta frequencies following the intervention. Positive values indicate excitatory coupling, while negative values indicate inhibitory coupling. Error bars represent SEM.

### 3.3 Changes in Cortical Activity Ratio (CAR)

We then tested for any intervention-induced changes in cortical activity for the 2 tasks related to motor execution. A 2 (time) x 2 (task) x 4 (region) repeated measures ANOVA found a significant Time * Task (*F*[1,7] = 8.03, *p* = 0.025, η_p_^2^=0.53) and Time * Region (*F*[3,21] = 4.64, *p* = 0.012, η_p_^2^=0.40) interaction. Post hoc paired t-tests found that following the intervention there was a significant decrease in the cortical activation in contralesional M1/S1 (*p* = 0.042) during hand opening (See Figure 3A). For the simultaneous lifting and opening condition, a significant increase in cortical activation in ipsilesional M1/S1 (*p* = 0.025) was observed (See Figure 3B).

**Figure 3.**
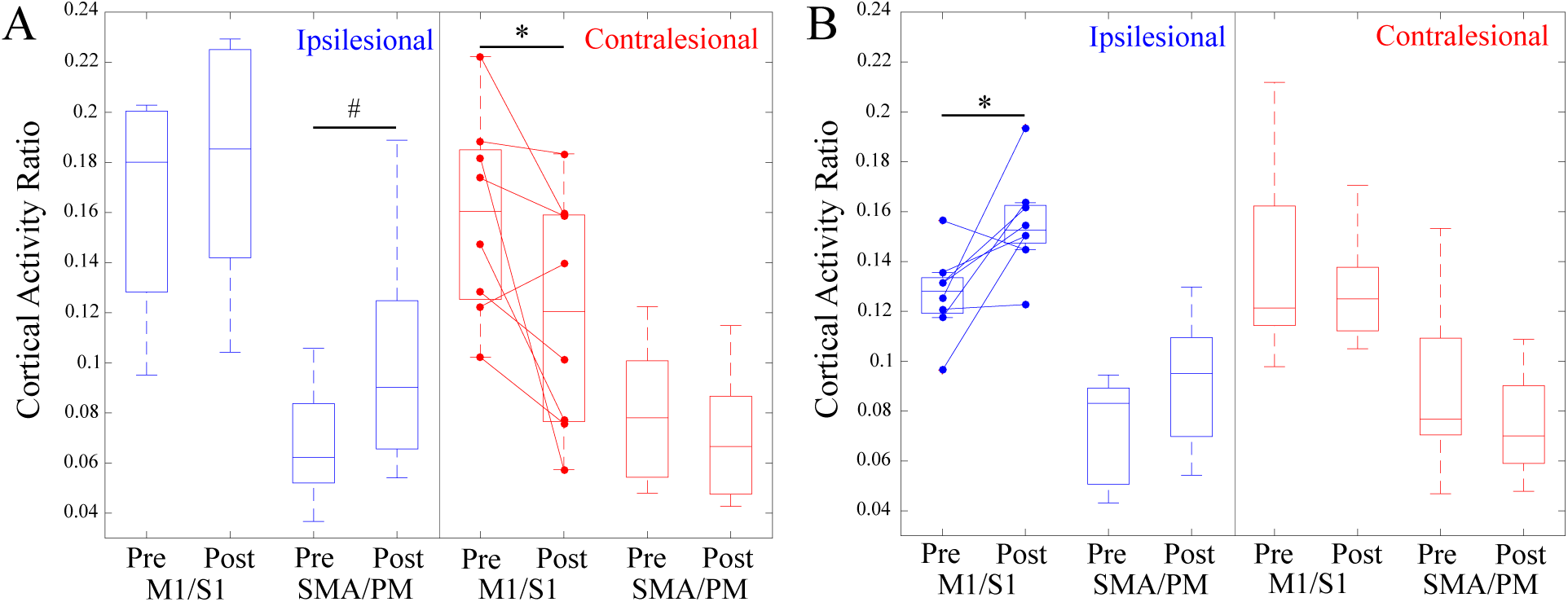
Box plots depicting cortical activity ratio Pre/Post Intervention for (A) hand opening on the table and (B) hand opening while lifting. Regions include combined primary motor cortex and primary sensory cortex (M1/S1) and combined supplementary motor area and premotor area (SMA/PM) for both the ipsilesional (Blue) and contralesional (Red) hemispheres. Participants demonstrated a decrease in contralesional primary sensorimotor cortex activity for hand opening following the intervention (left) and an increase in ipsilesional primary sensorimotor cortex activity for hand opening while lifting following the intervention (right). * indicates p < 0.05.

## 4. Discussion

The current study quantified changes in both cortico-cortico coupling during motor preparation and cortical topography at motor execution related to paretic hand opening with and without coordination of shoulder abduction following a device-assisted intervention. Overall, we observed that intervention-induced changes in coupling during motor preparation accompanied similar changes in cortical activity at motor execution. For hand opening, this was characterized by a reduction in coupling with contralesional M1 during motor preparation and similarly decreased activity in contralesional M1 at motor execution. For hand opening while lifting, this was characterized by a change in coupling within ipsilesional motor cortex during motor preparation followed by an increase in activity in ipsilesional motor cortex at execution.

### 4.1 Intervention-Induced Changes Related to Paretic Hand Opening

As we reported previously, participants demonstrated decreased contralesional primary sensorimotor activity related to paretic hand opening following the intervention. In this study, we further demonstrated that following the intervention, these individuals demonstrated a reduction in coupling from ipsilesional M1 to contralesional M1 within gamma frequencies.

In healthy controls, gamma band power increases in contralateral primary sensorimotor cortex just before the onset and during movement^37–42^. This increase in gamma power has been shown to facilitate movement in studies using transcranial alternation current stimulation (tACS) to artificially increase gamma power over the contralateral primary sensorimotor cortex^43–45^. Typically, gamma is associated with local intracortical processing, particularly within GABAergic interneuronal circuits^14,15^, with increases in gamma power related to movement associated with decreases in GABA_A_^44^. However, gamma synchronization is notable usually confined to a small area in the contralateral primary sensorimotor cortex in healthy controls^37–40,46^, rather than bilateral increases in gamma during movement as observed here prior to the intervention.

Before the intervention, individuals with severe chronic stroke showed positive gamma coupling (as demonstrated by the positive coupling shown in Figure 2B) from ipsilesional M1 to contralesional M1. This abnormal initial positive gamma coupling between ipsilesional and contralesional M1 may indicate abnormal intercortical communication between the two hemispheres (via callosal or subcortical means^47^) and may give rise to the initial increased contralesional activity in these individuals. This is supported by previous findings in stroke showing weaker GABAergic inhibition from the ipsilesional to the contralesional hemisphere following stroke^48–51^, and that this imbalance is associated with greater abnormal contralesional activity during paretic hand movements^52^.

Following the intervention, we found that the abnormal interhemispheric gamma coupling decreased in 6 out of 8 participants (see Figure 1B), which supports the intervention-induced reduction in contralesional activity in the primary sensorimotor cortex (see Figure 3A). The intervention-induced decrease in positive gamma coupling between ipsilesional and contralesional M1 may underlie the subsequent decrease in CAR in contralesional primary sensorimotor cortex and reflect a return to a more normal state as observed in healthy controls.

### 4.2 Intervention-Induced Changes Related to Paretic Hand Opening While Lifting

For the simultaneous lifting and opening task, we observed altered coupling within ipsilesional M1 in which the intervention induced a shift from positive to negative coupling between gamma and beta within ipsilesional M1 during motor preparation (see Figure 2B). This change was accompanied by a subsequent increase in ipsilesional primary sensorimotor cortex activity at motor execution following the intervention. The observed intervention-induced change in cross-frequency coupling is particularly relevant since gamma and beta power are typically inversely related in healthy controls during movement^23^. Whereas gamma tends to increase in power prior to and during the onset of movement, beta decreases in power^38,53^. This decrease in beta power during motor preparation in healthy controls has been linked with the reduction of inhibition in M1 just prior to movement to release the motor command^12,54^. Consequently, movements executed during elevated beta synchrony are slower^55^, and increasing beta power using tACS has been shown to impair movement in healthy controls^43,56^. Importantly, stroke individuals show less decreases in beta during movement compared to controls^16^, and persistence of inhibition in ipsilesional M1 during motor preparation^57^. Therefore, the shift from positive to negative coupling between gamma and beta in ipsilesional M1 following the intervention may reflect a return to a more typical pattern seen in healthy controls.

Prior to the intervention, individuals with severe chronic stroke demonstrated increased gamma power associated with increased beta power in ipsilesional M1 (as demonstrated by positive coupling between gamma and beta power in 6/8 participants, see Figure 2B). This positive coupling shifted to negative after the intervention, where increases in gamma power were associated with decreases in beta power. Importantly, decreases in beta power are inversely related to BOLD activity in the sensorimotor cortex, suggesting some interplay or association between these two physiological processes^58,59^. Given beta’s role in descending layer V pyramidal neurons^60,61^, the observed intervention-induced increase in CAR may reflect increased drive and use of remaining descending ipsilesional motor resources during the lifting and opening condition following the intervention, rather than purely an intracortical change. This is significant since reliance on descending tracts from the contralesional hemisphere has been linked with synergy-induced impairments^62,63^, particularly during tasks involving lifting at the shoulder^26,64^, as examined here. Meanwhile, increased use of descending corticospinal tract from the ipsilesional hemisphere has been shown to be crucial for improved function following stroke^65–67^, especially for independent control of multijoint movements of the upper extreminty^68^. Although contralesional corticobulbar pathways can support more proximal paretic arm movements such as reaching^69^, they do not offer sufficient control of independent joints during multijoint movements^70,71^. Thus, the ability to maintain ipsilesional recruitment during combined shoulder-hand tasks is critical for potential functional improvement since ipsilesional corticospinal tract, unlike contralesional corticobulbar tracts, has more specific branching in the spinal cord that allows for independent control of the different parts of the arm^71–74^.

### 4.3 Limitations

One of the major limitations of the current study is the small sample size (N=8). However, these participants are fairly homogeneous in that they are each severely impaired (FMA: 10-24; Chedoke McMaster: 3-4) with subcortical lesions impacting the posterior limb of the internal capsule and are in the chronic phase. The robotic setup during the EEG experiment also allowed for a well-controlled environment to investigate cortical activity related to the paretic arm and hand, thus minimizing potential session to session variability.

There are numerous methods to analyze connectivity besides DCM-IR when measuring EEG. Methods such as correlation or coherence between two time-series are the most commonly used metrics. However, unlike DCM-IR, these are limited to within-frequency relationships. Methods evaluating both within and cross-frequency relationships are rarer, including bicoherence, n:m phase synchrony, and multi-spectral phase coherence methods^75–78^. However, these methods either do not evaluate directionality, like correlations and coherence, or do not directly link to cortical regions. DCM-IR allowed us to evaluate both linear and nonlinear relationships between regions within a network, while also looking at directionality. However, DCM-IR only examines the temporal dynamics of power, but does not contain information of phase, which has been shown to play a potential role in cortical communication^18^. Lastly, DCM is limited by the models tested, and it is possible that there is a better model not considered here.

EEG is inherently limited in its spatial resolution, particularly in comparison to fMRI. However, using our high-density EEG setup, along with the sLORETA inverse calculation based on subject-specific boundary element models created with individual’s anatomical MRI, we have demonstrated a resolution of roughly 5 mm^79,80^. This is suitable for distinguishing primary sensorimotor cortex and secondary motor areas as investigated here in our topographical analysis and allowed us to investigate multi-joint movements which would have been impractical inside an MRI scanner. Meanwhile, the source reconstruction in DCM is based on an equivalent current dipole (ECD) method using the coordinates chosen based on our previous cortical activity findings in combination with standardized locations from the literature. Although these dipoles capture a smoothed portion of the cortex, it is possible that additional activity is not captured within the 5 regions investigated. It is worth noting that these 2 independent inverse approaches (sLORETA and ECD) yielded complementary results in the regions investigated.

## 5. Conclusion

The current study demonstrates that intervention-induced changes in coupling both within and between motor regions during motor preparation in movement-related frequencies complement changes in cortical activity at motor execution related to both the hand and arm. Together, these findings suggest that changes in cortico-cortico interactions during motor preparation may lead to corresponding changes in focal cortical activity.

## Conflict of Interest

None of the authors have potential conflicts of interest to be disclosed.

## Acknowledgements

The authors want to acknowledge Justin Drogos and Carolina Carmona for assistance with the intervention and Dr. Meriel Owen for MRI collection.

## Funding

This work was supported by an HHS grant 90IF0090-01-00 (formerly DOE NIDRR H133G120287), a NICHD 2RO1H-D039343 grant, and an award from the American Heart Association co-funded by the William Randolph Hearst Foundation 18PRE34030432.

## Ethical Approval

All procedures performed in studies involving human participants were in accordance with the ethical standards of the institutional and/or national research committee with the 1964 Helsinki declaration and its later amendments or comparable ethical standards. All participants provided written informed consent.

## Author Contributions

KBW, JPAD, and JY designed the study. KBW and JY collected the data. KBW conducted the analysis. KBW, JPAD, and JY wrote the manuscript.

## Competing Interests

The authors declare no competing interests.

## Data Availability

Data is available upon request.

## Notes

#### Summary of Updates

Alterations to abstract and introduction as well as slight aesthetic changes to figures.

