## Supplementary Material for "Cortical changes in motor preparation facilitate changes in motor execution following an intervention in severe chronic hemiparetic stroke"

**Supplementary Figure 1.** Models tested for DCM analysis. Models 1-6 allow only linear intrinsic connections and both nonlinear and linear extrinsic connections, while Models 7-12 allow both nonlinear and linear intrinsic and extrinsic connections. Individual models differ in interhemispheric connections allowed between M1 and PM regions. Dashed lines indicate only linear connections allowed, while solid lines indicate both linear and nonlinear connections allowed. The left side is the lesioned side.

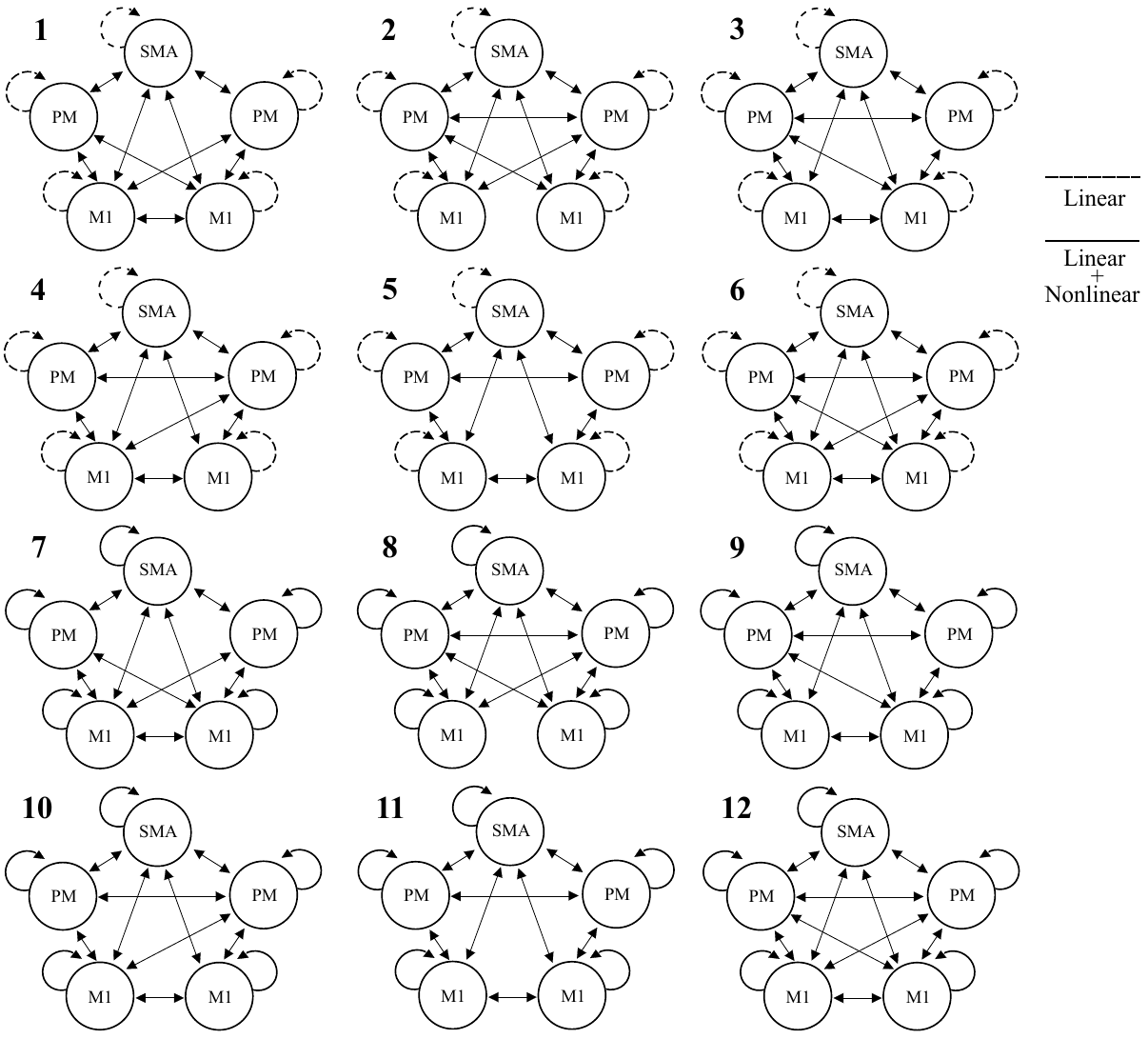

**Supplementary Table 1. Exceedance probabilities for each model**

| Models | Open | Lift + Open |
| --- | --- | --- |
| 1 | 0.0018 | 0.0022 |
| 2 | 0.002 | 0.0024 |
| 3 | 0.0028 | 0.0017 |
| 4 | 0.0023 | 0.0022 |
| 5 | 0.0021 | 0.0013 |
| 6 | 0.0022 | 0.002 |
| 7 | 0.0056 | 0.0029 |
| 8 | 0.0021 | 0.0037 |
| 9 | 0.0019 | 0.0026 |
| 10 | 0.0021 | 0.0024 |
| 11 | 0.0025 | 0.0018 |
| 12 | 0.9726 | 0.9748 |

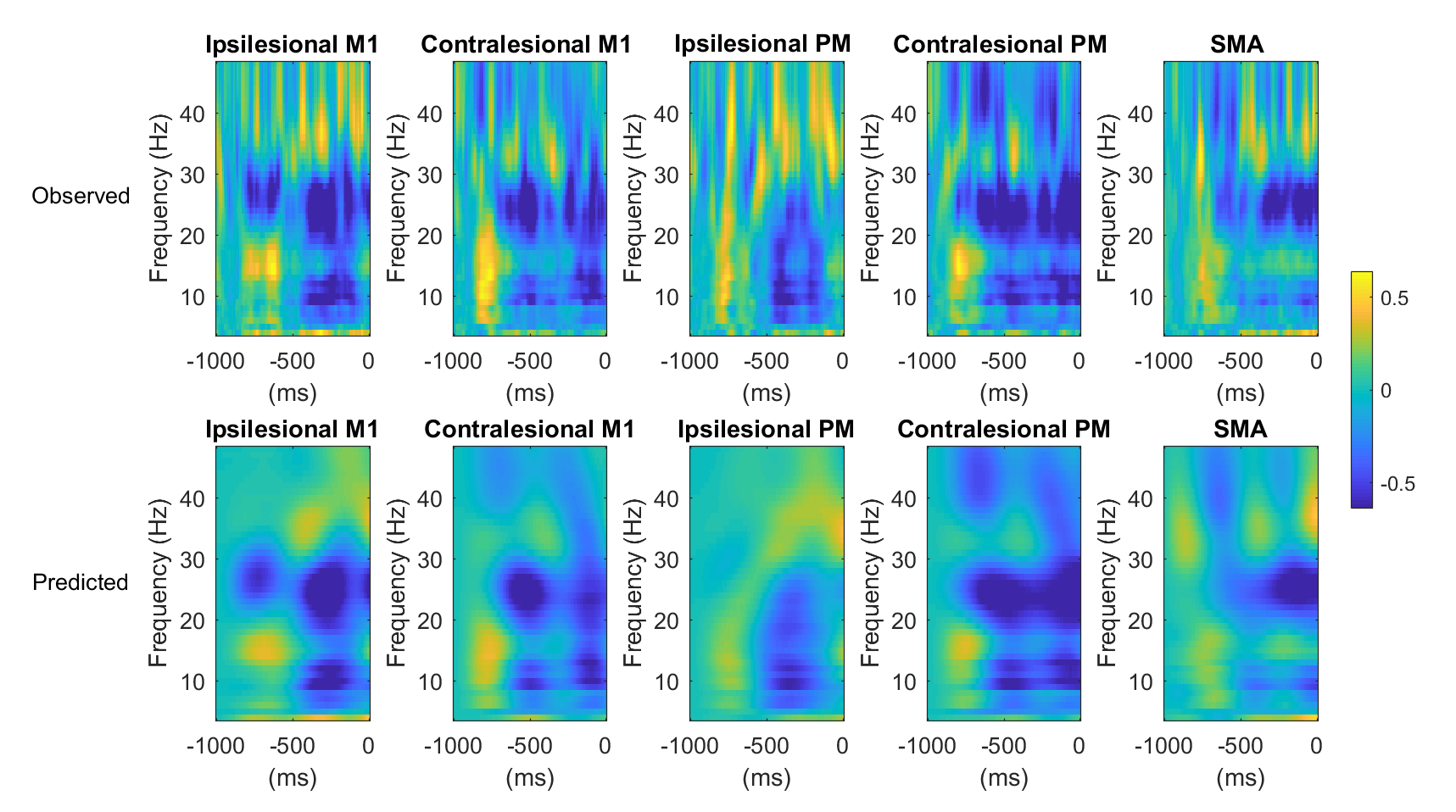

**Supplementary Figure 2.** The observed (top) and model-predicted (bottom) spectrograms for each region for one participant using the winning model (Model 12). Yellow indicates an increase in power compared to baseline and blue indicates a decrease in power compared to baseline. 0 ms indicates movement onset. Overall, the model explained ~85% of the original spectral variance for each condition.
